# Role of *Trypanosoma cruzi* nucleoside diphosphate kinase 1 in DNA-damage responses

**DOI:** 10.1101/830257

**Authors:** Chantal Reigada, Melisa Sayé, Fabio Di Girolamo, Edward A. Valera-Vera, Claudio A. Pereira, Mariana R. Miranda

## Abstract

NME23/NDPK proteins are well conserved proteins found in all living organism. Besides their catalytic activity of nucleoside diphosphate kinase (NDPK) they are considered multifunctional, which were first characterized as non-metastatic proteins in mammalian cells. Later, increasing evidences placed NME/NDPK as proteins involved in DNA stability such as gene regulation and DNA-repair. TcNDPK1 is the canonical NDPK isoform present in the parasite *Trypanosoma cruzi*, orthologous to NME23-H1/H2 which has been shown to have in vitro nuclease activity and DNA-binding properties. In the present study we investigate the role of TcNDPK1 in DNA-damage responses using heterologous gene expression systems and over-expression in epimastigote cells. We found that different strains of bacteria, WT and *ndk*-mutants, expressing the enzyme decreased about 5 fold and 18 fold the spontaneous mutation rate, respectively. In addition, yeasts lacking the endogenous gene *YNK1* (*YNK1-*) and expressing TcNDPK1, were significantly more resistant to different concentrations of hydrogen peroxide and were less sensible to UV radiation than controls. Parasites over-expressing TcNDPK1 were able to withstand different genotoxic stresses caused by hydrogen peroxide, phleomycin and hidroxyurea. In addition, under oxidative damage, TcNDPK1 over-expressing parasites presented lesser genomic damage and augmented levels of poly(ADP)ribose and poly(ADP)ribose polymerase, an enzyme involved in DNA repair. These results strongly suggest that TcNDPK1 is involved in the maintenance of parasite genomic-DNA integrity, thus, giving rise to a novel function.

## Introduction

Nucleoside diphosphate kinases (NDPK) are ubiquitous and well conserved enzymes whose canonical and first discovered function is to maintain the intracellular di- and tri-phosphate nucleotide homeostasis by catalysing their interconversion [1]. In addition, NDPK proteins are considered multifunctional enzymes since they have been shown to be involved in diverse physiological and pathological processes. Human NDPKs, comprised in the NME (NME23/NDPK) family, are by far the best studied NDPKs because of their prime role as metastasis suppressors [2]. The different isoforms have been found to be distributed all along the intracellular compartments, from cytoplasm to mitochondria, membrane and nucleus [3, 4]. Nuclear functions of NME proteins are currently being investigated in order to better understand the mechanisms underlying its non-metastatic function. NME-H1 and NMEH-2 have been identified as potential transcription factors that regulate gene transcription through their DNA-binding activities [5–9]. In addition, NME-H1, NME-H5, NME-H7, and NME-H8 also exhibit a 3′– 5′ exonuclease activity, suggesting roles in DNA proofreading and repair [10]. In fact, there are increasing studies confirming a possible role for NME-H1 in the repair of DNA damages induced by UV and gamma radiation, bleomycin, cisplatin or etoposide in different cell lines [10, 11]which is associated with an increased nuclear localization of the proteins.

The DNA-related functions of NDPK proteins have also been investigated in different organisms other than mammals, such as bacteria, plants, yeasts and protozoan parasites. A plenty of reports commit NDPKs to DNA-processing activities because of their interaction with DNA and their capacity to cleave it [12–17]. Furthermore, accordingly to what was reported for human NME proteins, *Deinococcus radiodurans* Ndk is up-regulated during gamma radiation stress [18] and the yeast homologue YNK1 is required for repair of UV radiation- and etoposide-induced DNA damage [19]. *Trypanosoma cruzi* is the protozoan parasite that causes Chagas disease. We have previously identified and characterized the *T. cruzi* NM23-H1 and NM23-H2 homologue, TcNDPK1, which is the only canonical NDPK present in the parasite [20]. TcNDPK1 presents 54% and 64% identity with NM23-H1 and NM23-H2 respectively and share some common functional features of most of the canonical NDPKs since, besides to the kinase activity, TcNDPK1 possesses unspecific in vitro nuclease activity [12], DNA-binding properties (Duhagon et al, unpublished) and cytosolic and peri-nuclear localization [21]. In addition, TcNDPK1 gene expression is modified in response to gamma radiation [22].To date, there are few reports involving classical NDPKs of trypasomatids in novel cellular processes; *T. brucei* NDPK have been localized to the nucleus without any other functional characterization [23] and *Leishmania amazonensis* homologue was mainly described as a protein implicated in the cell infection by preventing ATP-mediated cytolysis of macrophages [24]. Thus, in the present study we took the first steps to investigate the role of TcNDPK1 in DNA repair, finding that TcNDPK1 is involved in the maintenance of parasite genome integrity.

## Materials and Methods

### Parasites, yeasts and bacteria strains

Epimastigotes of *T. cruzi*, Y strains (DTUII), were used in all the experiments. Parasites were cultured at 28 °C in plastic flasks (25 cm^2^) containing 5 mL of LIT medium (started with 10^6^ cells/ml) supplemented with 10% fetal calf serum, 100 U/mL penicillin, and 100 mg/mL streptomycin [25]. The parasites were subcultured with passages each 7 days. Genetic constructions and transgenic parasites were obtained as previously reported [21]. Briefly, epimastigotes were transfected with a pTREX-TcNDPK1 (*TcNDPK1*-TritrypDB: TcCLB.508707.200) and a pTREX-GFP vector previously generated [21, 26, 27], cloned and maintained in LIT medium supplemented with G418 500 µg/ml. Expression of TcNDPK1 was confirmed by enzymatic activity.

The mutant YNK1^−^ *Saccharomyces cerevisiae* strain corresponds to the Euroscarf collection (BY4741 MATa; *his3Δ1*; *leu2Δ0*; *met15Δ0; ura3Δ0, YKL067W::KanMX*), kindly provided by Dr. Paula Portela (FCEyN, UBA). Yeasts were transformed as previously described [28] with an empty p416 GPD plasmid or a p416-TcNDPK1 plasmid constructed by the insertion of TcNDKP1 gene from the pTREX-TcNDPK1 in the BamHI and XhoI recognition sites of p416 GPD. Transformed yeast were checked by PCR and grown in selective ura^−^ medium supplemented with histidine, leucine and methionine.

Bacteria used in the study correspond to *E. coli* BL21 (DE3) strain transformed with an empty pRSET-A and a pRSET-TcNDPK1 previously generated [20]; and *E. coli* K-12 strain, *ndk*^−^ mutant from the Keio collection [29], kindly provided by Dr. Hirotada Mori (Nara Institute of Science and Technology, Japan) transformed with a pJexpress-404 bearing the gene of TcNDPK1 cloned in the BamHI and XhoI recognition sites (pJexpress-TcNDPK1). Both plasmids possess ampicillin (AMP) resistance cassette and K-12 *ndk*^−^ mutant kanamycin (KAN) resistance cassette inserted in the *ndk* gene. Expression of TcNDPK1 was evaluated by standard protocols for his-tag protein purification and enzymatic activity.

All the constructions made for the study were checked by sequencing.

### Western blots

5 10^7^/ml exponentially growing epimastigotes were treated in LIT medium with H_2_O_2_ 3 mM and aliquots containing 10^7^ parasites were taken at different times (0, 5, 10, 20 and 30 min), immediately centrifuged and lysed in 15 µl cracking buffer. Samples were cracked at 65°C to avoid cleavage of PARP and totally loaded in a 15% acrylamide gel. BSA (66 KDa) and ovalbumin (45 KDa) were used as molecular weight markers. The proteins were transferred to a PVDF membrane, blocked for an hour with 5% non-fatty milk in T-PBS buffer (0.05% Tween20, PBS 1x), incubated over night with rabbit anti-PARP antibodies (Santa Cruz Biotechnology) diluted 1:500 or anti-PAR reagent (MABE 1016, Millipore) diluted 1:500 in blocking buffer and finally incubated with HRP-conjugated anti-rabbit (Vector laboratories) diluted 1:5000 in blocking buffer. The membranes were revealed with ECL reagent (Pierce). Parasites viability was assessed by direct light microscope observations and MTS colorimetric assays (CellTiter Aqueous One Solution Cell Proliferation Assay -Promega); they maintained shape and motility at 5, 10 and 20 min post-treatment, at 30 min motility was affected and viability decreased about 25%.

### Immunofluorescence

Immunofluorescences were carried out as previously reported, using TcNDPK1 over-expressing parasites (N1 parasites) and the same batch of polyclonal mouse anti-TcNDPK1 antibodies previously used by Pereira et al. [21]. Briefly, parasites treated with H_2_O_2_ 0.3 mM for 1 h or with phleomicyn 150 µM for 4 h in LIT medium or without treatment (control) were settled on poly-L-lysine coated coverslips, fixed at room temperature for 20 min with 4% formaldehyde in PBS and permeabilized with cold methanol. After rehydration, the samples were blocked 10 min with 1% PBS-BSA and incubated 45 min with mouse anti-N1 serum diluted 1:50. Then parasites were incubated 30 min with anti-mouse-DyLight 488, diluted 1:500 in blocking buffer. Cells were mounted with Vectashield with DAPI (Vector Laboratories) and observed in an Olympus BX60 fluorescence microscope. Images were recorded with an Olympus XM10 camera.

### Nucleoside diphosphate kinase activity

Parasites were collected, counted in a Neubauer chamber, washed with PBS and resuspended in 100 mM Tris–HCl buffer pH 7.2. Protein extracts were obtained by 7 freezing and thawing cycles in liquid N_2_ followed by centrifugation. The activity was measured in the supernatant by measuring NADH oxidation at A_340nm_ as previously reported [20].

### Genomic samples and electrophoresis

5 10^7^/ml parasites were treated with H_2_O_2_ 3 mM in LIT medium and 1 ml-samples were collected at different times (0, 5, 10, 20 and 30 min), immediately centrifuged and resuspended in 0.3 ml of lysis buffer (10 mM Tris buffer pH 7.5, 100 mM EDTA, 0.1 % SDS). These solutions were incubated with RNAse 30 min at 37 °C and then extracted with 0.3 ml of phenol. The aqueous phases were extracted with 0.3 ml of chloroform-isoamyl solution and genomic DNA was precipitated with isopropanol and recovered by centrifugation. Pellets were resuspended in 15 µl of TE buffer and quantified in a NanoDrop Nucleic Acid Quantification (Thermo Fisher). 200 ng of genomic DNA were loaded in an ethidium bromide-stained 1% agarose gel, electrophoresed at 50V for 2 h and visualized in a UVC device.

### Genotoxic stresses

7.5 10^6^ exponentially growing epimastigotes were incubated in a 24 wells plate with different concentration of H_2_O_2_ (0, 50, 100 and 125 µM) in 300 µl of PBS at 28 °c for 2h. Then, 1.2 ml of fresh LIT medium was added (5 10^6^ parasites/ml final) and parasites were incubated for other 96 h as previously reported [30]. For phleomycin and HU treatments, same density of parasites were incubated in 500 µl of LIT medium with different concentrations of each genotoxic agents (Phleo: 0, 60 and 100 µg/ml; HU: 0, 10 and 20 mM) for 96 h and 24 h respectively, as previously reported [31]. After treatment, culture growth was monitored by parasite counting in a Neubauer chamber using an Olympus BX60 microscope.

In the case of *S. cerevisiae*, experiments started from an overnight culture. H_2_O_2_ treatment was carried out by incubating 0.4 OD in 0.2 ml of PBS with different concentration of H_2_O_2_ (0, 5, 10, 25 and 50 mM) for 30 min. Then 2.8 ml of selective medium was added and incubated in agitation at 30°C for 24 h. Yeast growth was measure by recording A_600nm_. For UV treatment, yeasts were diluted to 1 OD/ml (approximately, 10^7^ /ml) and 50 µl of a 10^−4^ dilution was loaded into selective ura^−^ agar in a 6 cm petri plate and subjected to UVC radiation in a UVP gel Imaging and Documentation device (Bio-Rad) for 15 and 30 sec. Then, plates were incubated at 30°C for 96 h and colonies were counted. Survival percentages were obtained by taking as 100% the number of colonies without radiation.

### Rifampicin mutation frequency

For K-12 strains, 12–20 cultures were started from a single colony in 3 ml 2YP-lactose broth (added with 0.5 mM lactose to allow basal transgene expression) and grown over night at 30°C to avoid over-saturation with the corresponding antibiotics (K12 ndk^−^: 30 µg/ml KAN; K-12 ndk^−^/TcNDPK1: 30 µg/ml KAN and 100 µg/ml AMP). The number of Rif^R^ mutants in each culture was determined by plating 1ml of undiluted cultures on LB-Rif plates (100 µg/ml Rif, 30 µg/ml KAN and 100 µg/ml AMP when corresponding), and the total number of viable bacteria by plating 100 µl of a 10^−6^ dilution on LB plates. Frequencies were obtained by dividing the number of Rif^R^ mutants per total number of bacteria in each culture. Then, the median from each set of cultures was recorded. Sporadic jackpot cultures were removed from the analysis.

For BL21 strain, a similar approach was carried out, but 2YP-100 µg/ml AMP broth was used instead. Expression of transgene was due to the leaky expression of the pRSET system [20].

### Data analysis

All the experiments were made at least in triplicate (replicates) and results presented here are representative of three independent assays. When corresponding, data was analysed by a two-way ANNOVA corrected by Sidack’s multiple comparison test with 95% confidence interval, using GraphPad Prism 6.01 for Windows.

Densitometries were carried out with the free available software ImageJ. Densities were normalized with bands of the corresponding Ponceau S red-stained membranes (loading control) or maxicircle bands and then referred to 0 min of N1 parasites.

## Results

### 1 Bacteria expressing TcNDPK1 have a lower spontaneous mutation frequency

In order to determine if TcNDPK1 is involved in DNA-repair processes we first measure its effect on bacterial spontaneous mutation frequency, assessed by determining the rifampicin resistant (Rif^R^) mutation frequency. We transformed *E. coli* K12 mutants lacking the endogenous NDPK gene (K12 ndk-) with a pJexpress plasmid bearing the TcNDPK1 gene. The expression of TcNDPK1 (K12 ndk1-/TcNDPK1) reduced the mutation frequency around 18 fold than the control without the plasmid (K12 ndk-), while not reaching the frequency of K12 WT bacteria which was 43 fold lower than mutants and 2.3 fold lower than K12 ndk1-/TcNDPK1. In addition, the expression of the TcNDPK1 gene in *E. coli* BL21 bacteria using the pRSET vector produced a decrease of 5 fold the spontaneous mutation frequency (Table 1).

**Table 1.** *Bacterial mutation frequency*. The Rif^R^ mutation frequency was calculated in *E. coli* K-12 bacterias, WT or mutants lacking the endogenous *ndk* gene, and in *E. coli* BL21 DE3, expressing or not TcNDPK1. The median of 12-20 independent assays is reported; in brackets are the ranges of frequencies obtained in each experiment. a, compared to WT; b, compared to BL21 pRSET.

| Strain | No. Rif<sup>R</sup> per 10<sup>8</sup> cells | Decreased folds |
| --- | --- | --- |
| *K-12 wild-type* | 0.914 (0.514-1.96) | 43.32<sup>a</sup> |
| *K-12 ndk<sup>-</sup>* | 39.6 (18.7-85.8) | 0 |
| *K-12 ndk<sup>-</sup>/TcNDPK1* | 2.15 (1.53-4.77) | 18.41<sup>a</sup> |
| *BL21 pRSET* | 1.85 (1.21-3.6) | 0 |
| *BL21 pRSET-TcNDPK1* | 0.352 (0.1-0.98) | 5.2<sup>b</sup> |

### 2 Yeasts expressing TcNDPK1 are more resistant to genotoxic agents

As was reported for *Saccharomyces cerevisiae YNK1* gene [19], the next step was to investigate the ability of TcNDPK1-expressing *S. cerevisiae* to survive to DNA damages induced by genotoxic agents. It is well established that hydrogen peroxide and UV radiation cause DNA damage [32, 33]; hydrogen peroxide by generating DNA oxidative damage and UV, by producing DNA cross-linking lesions. For this purpose, mutant yeasts lacking the endogenous NDPK gene (*YNK1*^−^) were transfected with the p416 vector bearing the *TcNDPK1* gene and treated with increasing concentrations of H_2_O_2_ or exposures to UVC radiation. Both yeast lines (N1 and control) had similar growth kinetics but yeast expressing the *T. cruzi* protein tolerated better the different stresses as is showed in figure 1 (Fig 1), being significantly more resistant to 5, 10, 25 and 50 mM hydrogen peroxide after 24 h of treatment and to 4.25 and 8.5 J/cm^2^ UV exposure than controls (Fig 1C).

**Figure 1.**
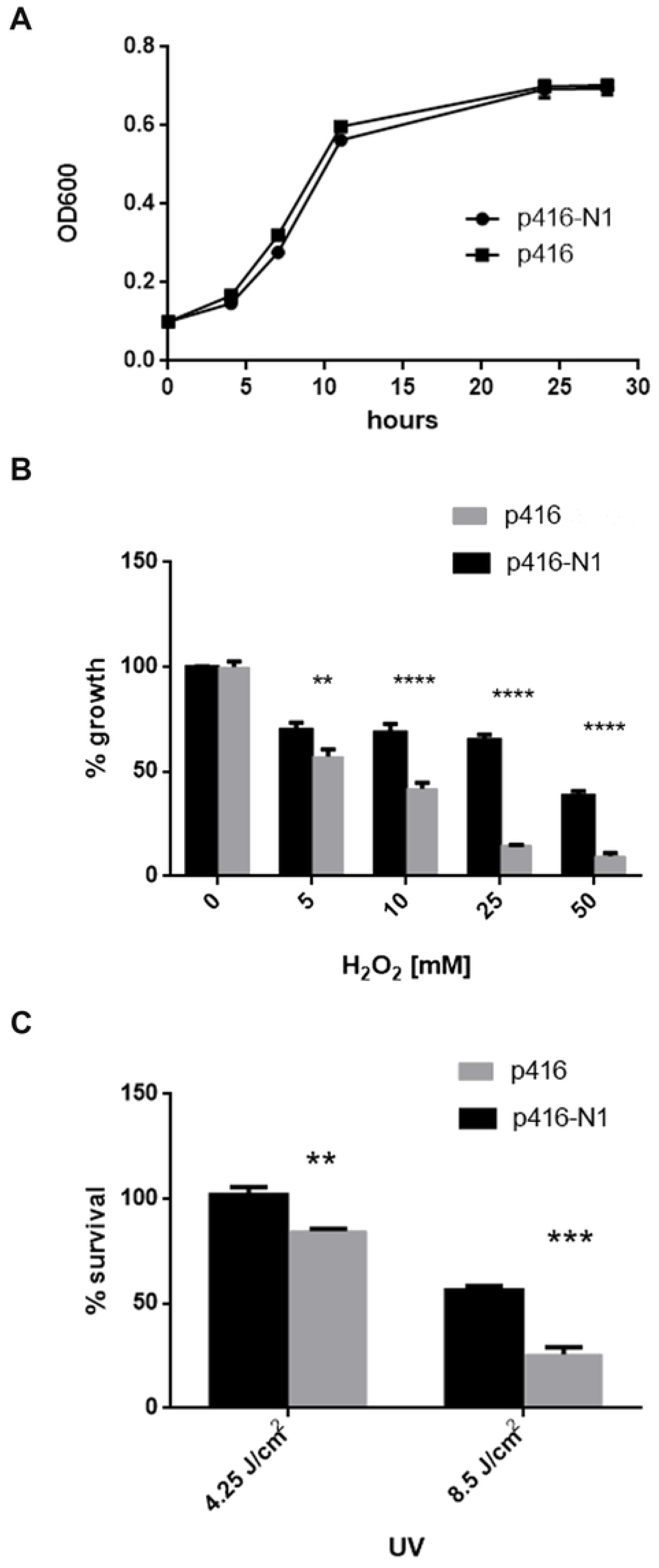
*Yeasts treatment with genotoxic agents*. Mutant yeasts lacking the endogenous *YNK1* but expressing the *TcNDPK1* gene were subjected to different genotoxic agents. A) Growth curve of transfected yeasts measured at A_600nm_. B) Percentage of yeast growth against H_2_O_2_ treatment. Yeasts were treated with different concentration of H_2_O_2_ and A_600nm_ was measured 24 h after treatment. Percentages were calculated taking H_2_O_2_ 0 mM as 100% C) Percentage of yeast survival against UVC radiation. Yeasts were loaded in petri dishes, subjected to 4.25 and 8.5 J/cm^2^ UVC radiation and incubated for 96 h. Percentages were calculated by counting colonies from radiated and non-radiated plates. ** p ≤ 0.01, *** p ≤ 0.001, **** p ≤ 0.0001

### 3 Epimastigotes over-expressing TcNDPK1 are more tolerant to genotoxic agents

In order to evaluate the role of TcNDPK1 in the ability of the parasites to withstand genotoxic stresses, we generated a TcNDPK1 over-expressing line of epimastigotes by transfected them with a pTREX-N1 plasmid (N1 parasites); control was obtained by expressing the GFP with the same plasmid (GFP parasites). N1 parasites presented increased NDPK activity, about eight fold respect to control; rates obtained were 79.1 TTP µmol/sec per 10^8^ parasites for N1 and 9.5 TTP µmol/sec per 10^8^ parasites for GFP. In addition, the transgenic lines had similar culture growth and duplication rates, reaching same densities at stationary phase (Fig 2A).

**Figure 2.**
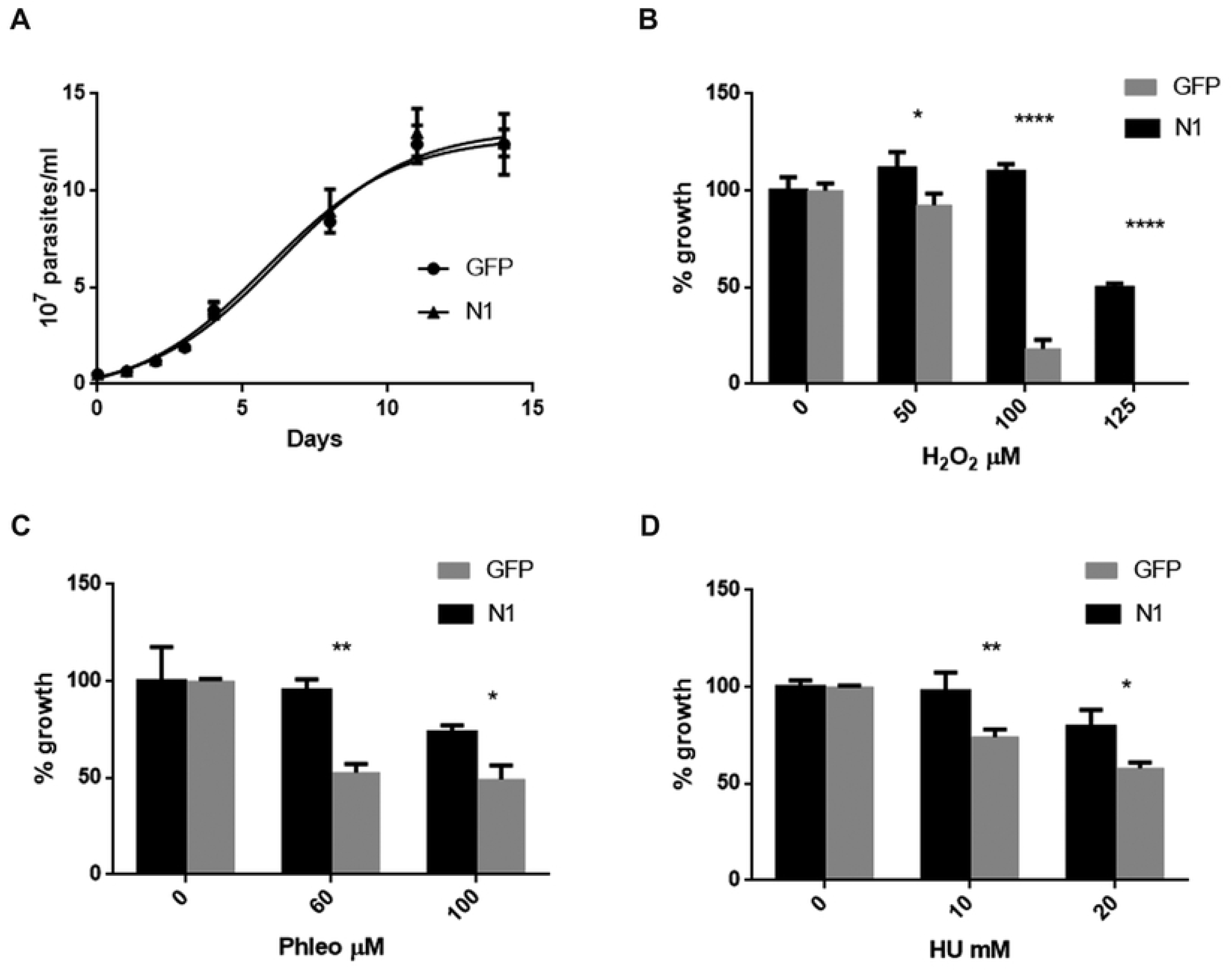
*Epimastigotes treatment with genotoxic agents*. Epimastigotes over-expressing the enzyme were cloned and A) growth was monitored by counting under microscope. Then, they were treated with different genotoxic agents: B) H_2_O_2_, C) phelomycin (Phleo), D) hydroxyurea (HU), and counted under microscope after 96 h (B and C) and 24 h (D) of treatment. Percentages were calculated taking 100% parasites without drug. GFP over expressing parasites were used as control. * p ≤ 0.05, ** p ≤ 0.01, **** p ≤ 0.0001

Both transgenic lines were treated with different concentrations of genotoxic agents that induce single and double DNA strand breaks and replication stress: H_2_O_2_, phleomycin (Phleo) and hydroxyurea (HU) as was reported by Gomes et al [31]. N1 parasites were more tolerant to all these DNA-damage inducing compounds as is shown in figure 2 (Fig 2). Markedly, at H_2_O_2_ 100 µM 100 % of N1 parasites survived to the treatment while GFP parasites grew only 20 %; in addition, H_2_O_2_ 125 µM was 100% lethal to GFP parasites and N1 population persisted reaching 50% of growth (Fig 2 B), measured 96 h after treatment. In contrast to H_2_O_2_, Phleo and HU stopped duplication maintaining parasites alive at the concentrations evaluated. Significant differences were observed at phleomycin 60 µg/ml and 100 µg/ml, reaching N1 parasites about 95% and 75% of growth respectively against 50% and 45% of control parasites (Fig 2C) after 96 h of treatment. Regarding HU, significant differences were also evidenced; N1 parasites were more resistant to 24 h of replication stress, since this population was able to endure about 24% more than GFP parasites at HU 10 mM and 20 mM (Fig 2D).

### 4 Epimastigotes over-expressing TcNDPK1 has less genomic DNA damage and enhanced activity of TcPARP

In previous works it has been demonstrated that treatment with H_2_O_2_ 0.03 to 3 mM generates DNA damage at short times as 10 min [34, 35]. To evaluate the genomic damage caused by H_2_O_2_ in the transgenic lines, we first incubated parasites with H_2_O_2_ 0.3 mM and the genomic DNA was extracted at different times (0, 15, 30 and 60 min) and visualized by agarose gel electrophoresis. At these conditions no DNA fragmentation was observed, so parasites were then exposed briefer times to 3 mM H_2_O_2_ (0, 5, 10, 20 and 30 min). Accordingly with the results obtained above, N1 parasites presented slightly less DNA damage than control parasites, as can be seen in figure 3 (Fig 3A and 3B).

**Figure 3.**
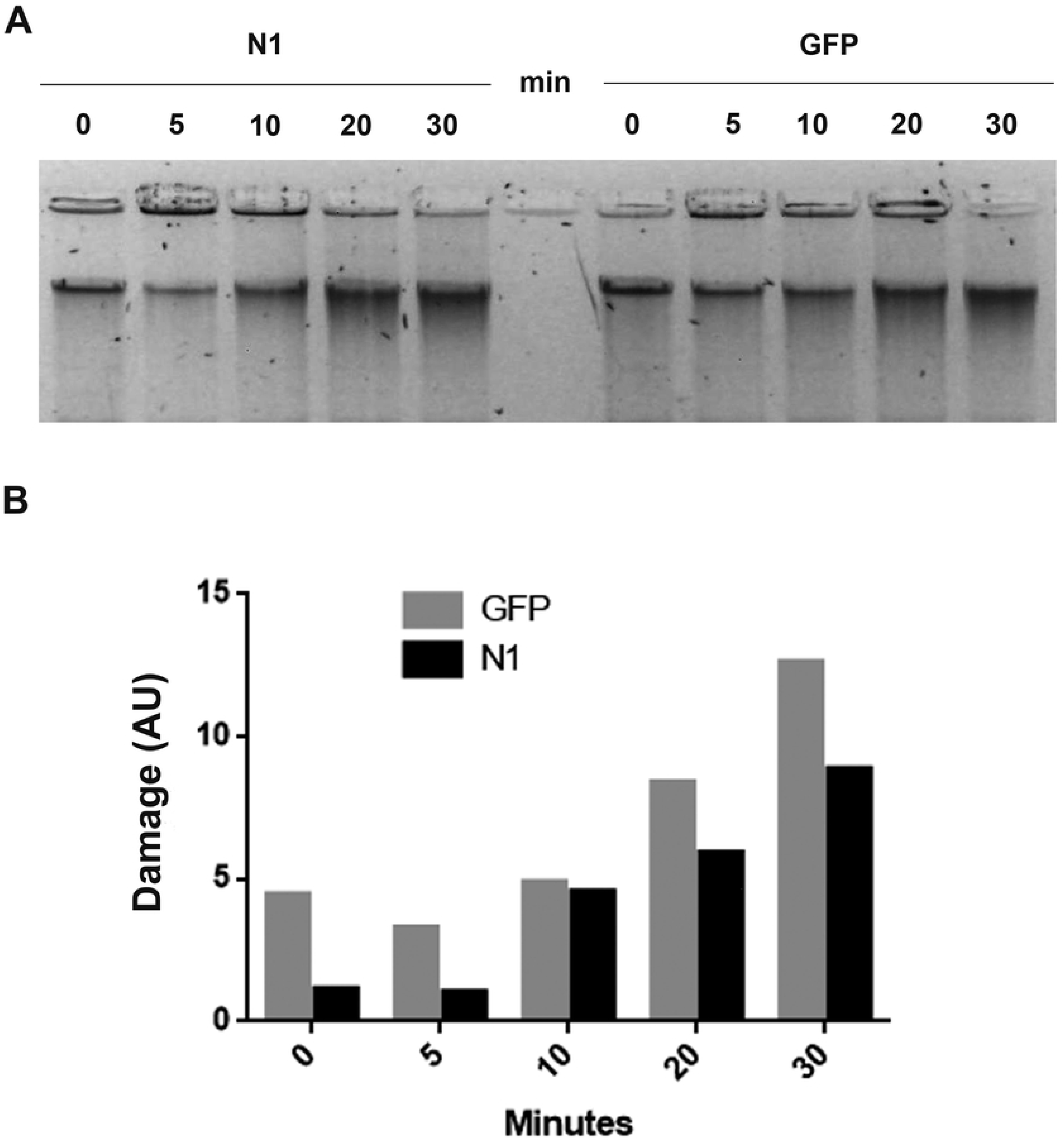
*Genomic DNA damage of epimastigotes*. Parasites were treated with H_2_O_2_ 3 mM and A) genomic DNA was isolated after 0, 5, 10, 20 and 30 min after treatment, loaded in an agarose gel and stained with ethidium bromide. B) Densitometry of A). GFP parasites were used as control.

Different DNA damage responses are activated under these genotoxic conditions. One DNA-break sensor enzyme that rapidly acts under DNA damage is the Poly(ADP-ribose) polymerase (PARP), which catalyses the PARylation of nuclear proteins. *T. cruzi* possesses only one PARP of 65 KDa (TcPARP) and it has been reported that at H_2_O_2_ 3 mM, activates and translocates to the nucleus of the parasites [34]. To further examine the activation of TcPARP in N1 and control populations, parasites were exposed to H_2_O_2_ 3 mM at different times (0, 5, 10, 20 and 30 min) and PARylation was evaluated by western blot (Fig 4A). In both parasites the level of modified proteins increased with damage until 20 min, but N1 parasites presented more intensity than control parasites and, remarkable, they maintained the elevated levels at 30 min against GFP parasites that showed an important decreased of about 46% at the same time (Fig 4B). The levels of TcPARP were also evaluated (Fig 4C) and in accordance with PARylation, the levels augmented with time exposure and in N1 parasites continued increasing at 20 min keeping them high at 30 min, whereas control parasites showed an abrupt decreased being almost undetectable at the mentioned times (Fig 4D). In addition, at 5 and 10 min, induction of expression was greater in N1 parasites, reaching about 6 to 10 fold respect to 0 min against 1.5 and 1.7 fold of GFP parasites (Fig 4D).

**Figure 4.**
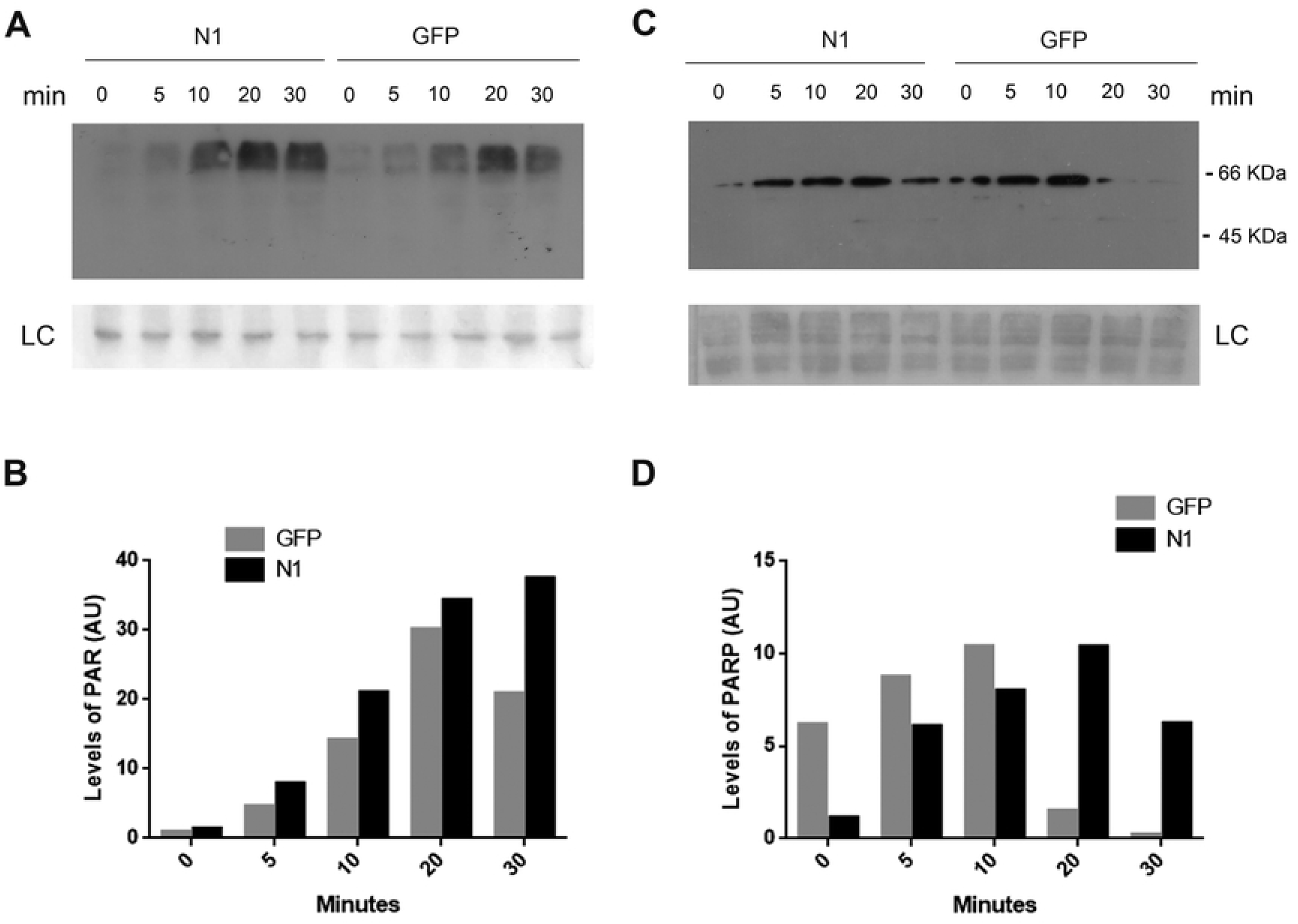
*TcPARP activity*. Parasites over-expressing TcNDPK1 were treated with H_2_O_2_ 3 mM and levels of A) PAR and C) TcPARP at 0, 5, 10, 20 and 30 min were measure by western blot. B) and D) are densitometries of A) and C) respectively. GFP parasites were used as control. LC: loading control, corresponding to Ponceau S red-stained membranes.

### 5 TcNDPK1 has nuclear localization

Previous experiments carried out in our laboratory threw the presence of NDPK activity in nuclear fractions (Miranda et al, unpublished) and, in addition, we determine that TcNDPK1 has a cytosolic localization, enriched mainly around the nucleus [20, 21] Furthermore, the orthologous gene of *TcNDPK1* in *T. brucei* codifies a NDPK that was reported as nuclear enzyme [23]. In order to evaluate in detail the subcellular localization of TcNDPK1, immunofluorescences were carried out in these transgenic parasites. As is shown in figure 5, in addition to be cytosolic, TcNDPK1 showed marked nuclear and peri-nuclear localization, very similar to its *T. brucei* counterpart (Fig 5, for details, please see [23]). In addition, phleomycin and H_2_O_2_ treatment produced a slight delocalization of TcNDPK1 out of the peri-nuclear region (Fig 5).

**Figure 5.**
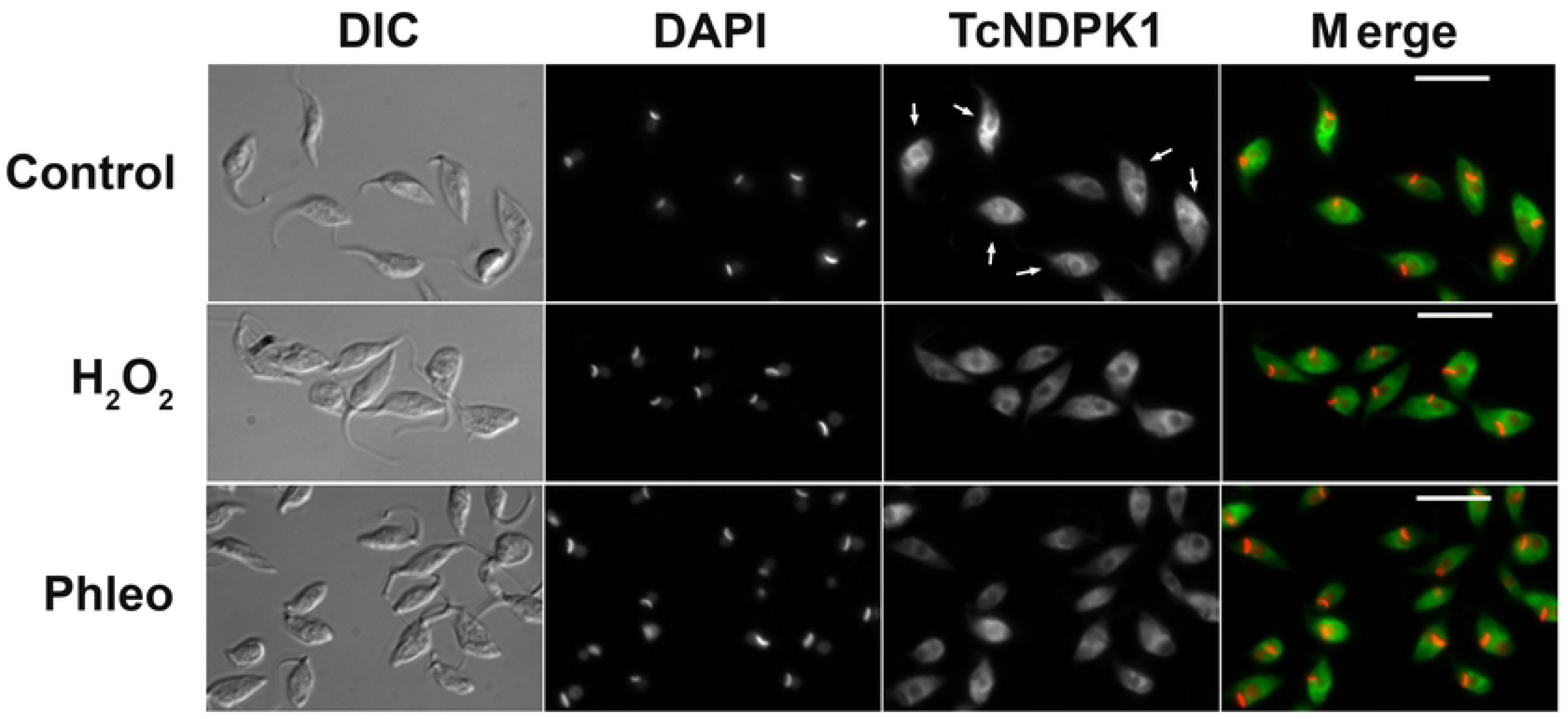
*TcNDPK1 localization*. Immunofluorescence of N1 parasites with anti-TcNDPK1 antibodies visualized with anti-mouse-DyLight 488. Parasites were no treated (control) or treated with H_2_O_2_ 0.3 mM for 1 h and with Phleo 150 µM for 4 h. Nucleus were stained with DAPI. Arrows point control parasites with a marked nuclear and peri-nuclear TcNDPK1 localization.

## Discussion

In the present study we have explored the role of TcNDPK1 in DNA-repair mechanisms providing the first evidences that involve this enzyme in this process.

One of the experimental strategies to identify possible proteins involved in DNA repair is by measuring the Rif^R^ mutation frequency of bacteria once incorporated the gene of interest. The heterologous expression of *TcNDPK1* in different bacterial genetic backgrounds produced a decreased in their spontaneous mutation frequency, thus suggesting a role of TcNDPK1 in the DNA repair machinery. Yeasts expressing TcNDPK1 were more tolerant to DNA-damage induced by H_2_O_2_ and UV radiation and over-expression of the enzyme in epimastigotes of *T. cruzi* generated parasites with improved ability to withstand to H_2_O_2_, phleomycin and HU –induced DNA lesions. Furthermore, less genomic DNA-damage could be observed in TcNDPK1 over-expressing parasites under H_2_O_2_ treatment. Altogether, these results offer new insights in the role of TcNDPK1 in DNA-damage responses. Supporting these evidences, Grynberg et al found that the expression of *TcNDPK1* is differentially regulated under gamma radiation exposure [22].

H_2_O_2_, UV radiation, Phleo and HU generate distinct forms of DNA damage. H_2_O_2_ produces oxidative stress generating single and double strand breaks (SSB and DSB, respectively), base modifications, apurinic/apirimidinic sites and DNA-protein crosslinks [32, 36]; UV radiation produces intra-strand cross linking and pyrimidine photoproducts [33]; Phleo is an antibiotic that binds to DNA and generates DSB [37–39] and HU is an antimitotic inhibitor of the enzyme ribonucleotide reductase, that causes replicative stress [40]. *T. cruzi* appears to have different ways to lead with DNA damage since most known DNA repair pathways are present in these organisms [36]. Increasing evidences indicate that strand breaks, especially DSB, are repaired by homology recombination (HR) in which TcRAD51 participates [31]. In addition, several enzymes involved in base excision repair (BER) and mismatch repair (MMR) have been characterized in *T. cruzi* which are responsible of repairing oxidative modifications of DNA bases, such as TcOGG1, TcMYH, TcMSH2 and TcMTH [41–44]. Probably, the genotoxic agents assessed in this study elicited different types of lesions so that more than one repair pathway could be overlapping. One interesting enzyme that appears to be shared by the different pathways is PARP. PARP acts as a DNA-damage sensor that promotes DNA repair at low levels of genotoxic stress, it activates by binding to DNA-SSB and –DSB and modifies other target protein with PAR chains [45]. *T. cruzi* possess one PARP, TcPARP, which, at the oxidative conditions assayed in the present study and under UVC radiation, translocates to the nucleus and increases the levels of PAR [34, 46]. In this scenario, to better understand a possible mechanism of action of TcNDPK1 in DNA-damage responses we evaluated the levels of PAR and TcPARP in the transgenic parasites. Under H_2_O_2_ DNA damage, PARylation was increased in N1 parasites as well as the induction of expression of TcPARP, indicating that the TcPARP activity might be enhanced due to changes in its expression. The most pronounced differences were observed at extended times of treatment where N1 parasites maintained elevated levels of PAR and TcPARP. These results suggest that, at least when dealing with DNA-oxidative damage, TcNDPK1 response may involve TcPARP. Interestingly, Vilchez Larrea et al showed that large amounts of PAR are produced in a H_2_O_2_ dose-dependent manner without changes in TcPARP levels [34], somehow similar to what was observed in GFP parasites but in contrast to N1 parasites. In addition, in *T. brucei* the lack of a functional TbPARP leads to an increase resistance against hydrogen peroxide and transgenic cells over-expressing TbPARP showed increased sensitivity to the same treatment [47]. Thus, differences could be explained in part by the action of TcNDPK1.

Increasing studies have confirmed a possible role for NN23-H1, one of the human orthologous of TcNDPK1, in DNA repair through the BER and nucleotide excision repair (NER) pathways and also in DSB repair [48]. Consistent with this role, NM23-H1 was reported to be peri-nuclear that rapidly translocate to sites of DNA damage in the nucleus [10, 11]The mechanism underlying nuclear translocation is poorly understood, since these enzymes lack a canonical nuclear localization signal, even though a passive diffusion is also expected due to their small molecular weight. Accordingly, we determine that TcNDPK1 possess cytosolic, peri-nuclear and nuclear localization, a pattern very similar to the one reported for *T. brueci* NDPK [23]. In addition, under oxidative damage and phleomycin treatment, the marked peri-nuclear localization seems to get lost. Stimuli-dependent translocation of NDPKs has also been reported in other organisms [49], suggesting that delocalization could be part of their regulation or mechanisms of action.

The present report constitutes an exploratory study of the role of TcNDPK1 in DNA-damage responses, and the results strongly suggest that TcNDPK1 is involved in genomic DNA maintenance. Further investigations will be needed to reveal in detail the extent to which TcNDPK1 participates in these processes. This knowledge will allow unravelling the intricate network of multiple functions this enzyme gathers.

## Acknowledgements

Special thanks to Dr. Hirotada Mori for kindly provide us the *NDK*^−^ mutant bacterias from the Keio Collection, Dr. Silvia Fernandez Villamil and Dr. Salome Vilchez Larrea for the anti-PAR reagent, Dr. Patricia Bustos for the anti-PARP antibodies and Dr. Paula Portela for the YNK1-mutant yeasts.

## Funding statement

This work was supported by Agencia Nacional de Promocion Científica y Tecnologica (ANPCyT). Grants: FONCYT PICT 2013-2218, 2015-0539 and 2018-01871. Global Challenges Research Fund (GCRF), Grant: MR/P027989/1. Consejo Nacional de Investigaciones Científicas y Tecnicas (CONICET) for salaries of researchers, fellowships and technicians: Claudio A. Pereira, Mariana R. Miranda, Melisa Saye, Chantal Reigada, Fabio Di Girolamo and Edward Valera-Vera.

